# Evaluating open LLMs for agentic analysis orchestration in a typical biomedical lab

**DOI:** 10.64898/2026.05.13.724985

**Authors:** Anton Nekrutenko

**Affiliations:** Department of Biochemistry and Molecular Biology, Penn State University, University Park, PA 16802

## Abstract

Agentic tools — software environments where a large language model plans, calls external tools, executes code, and iterates with minimal human intervention — will run a substantial share of routine biomedical data analysis within the next few years. However, per-call inference cost on frontier models is the bottleneck and can add up quickly. Here, we tested whether a free, locally-runnable open-weight model could take over the repetitive execution steps at frontier accuracy. We used Claude’s Opus to author plans of increasing detail for per-sample variant calling, and ran six 2026-release open-weight implementer LLMs against those plans on a set of desktop GPUs. qwen3.6:27b reproduced frontier accuracy on every plan and matched Opus cell-for-cell on a 36-cell error-injection matrix. A sub-$2,000 Jetson or Apple Mac Mini sufficed for the implementer side. The open-weight model landscape evolves on the order of months, so the specific implementer recommended here will be superseded; we provide the plans, harness, scoring code, and per-cell artifacts at https://github.com/nekrut/LLM-eval-paper as a framework for re-evaluating future models.

## Introduction

Frontier models from Anthropic, OpenAI, and Google write quality working code for data analysis tasks, but they cost cents to dollars per call, resulting in rapidly ballooning bills. It does not make sense asking Claude’s Opus to write a hundred-line bash script every time a few FASTQ files need to be aligned to reference wasting a token budget that should go to harder problems. Can we instead ask an expensive frontier model to write a recipe once, and then have a free, small open-weight model running on the lab’s own hardware turn that recipe into a working script every time after?

Prior LLM-in-bioinformatics evaluations fall into three categories. The earliest measured factual recall on genomics question-and-answer benchmarks: GeneGPT [1] gave a closed model live access to the NCBI web APIs; the 2025 GeneTuring re-evaluation [2] extended the design to 1,600 questions across sixteen model configurations spanning closed frontier systems and small biomedical models. A second category shifted to code generation on isolated tasks. BioCoder [3] graded 2,269 short bioinformatics functions against unit tests but restricted its open-source arm to pre-current-generation code-completion models. BioLLMBench [4] scored GPT-4, Gemini, and LLaMA on 24 tasks and found LLaMA frequently failed to emit runnable code; an evaluation of 104 Rosalind problems [5] placed GPT-3.5 ahead of GPT-4o and Llama-3-70B. A third category targets end-to-end pipelines and agentic systems — LLMs that plan, call tools, and revise their output in a loop. BixBench [6] grades closed models on more than 50 Dockerised computational-biology scenarios at ∼17 % open-answer accuracy; BioMaster [7] wraps a Plan/Task/Debug/Check loop around retrieval-augmented generation (RAG) for RNA-seq, ChIP-seq, scRNA-seq, and Hi-C analyses; BioAgents [8] fine-tunes small open-weight models for local execution. Two workflow-language studies generate Galaxy and Nextflow pipelines with DeepSeek-V3 [9] or rerank Galaxy workflows with Mistral-7B against GPT-4o [10].

Our study is different. We test whether a recipe authored once by a frontier model can be executed end-to-end by a free, locally-runnable open-weight implementer at frontier accuracy. Our goal is to characterize the recipe-implementer split on a real bioinformatics task — per-sample mtDNA variant calling on four paired-end Illumina samples — across the lab-scale hardware tier (MacBooks, a desktop with a gaming video card, a recycled workstation, and a small NVIDIA device). Briefly, claude-opus-4-7 authors a series of plans of increasing detail; six 2026-release open-weight implementers run those plans on a desktop GPU; the strongest implementer is then cross-platform validated and stress-tested with a programmatic error-injection.

To decide which open models to use we first need to survey the landscape of available models. For 2026 this landscape is summarized in Table 1 below.

**Table 1.**
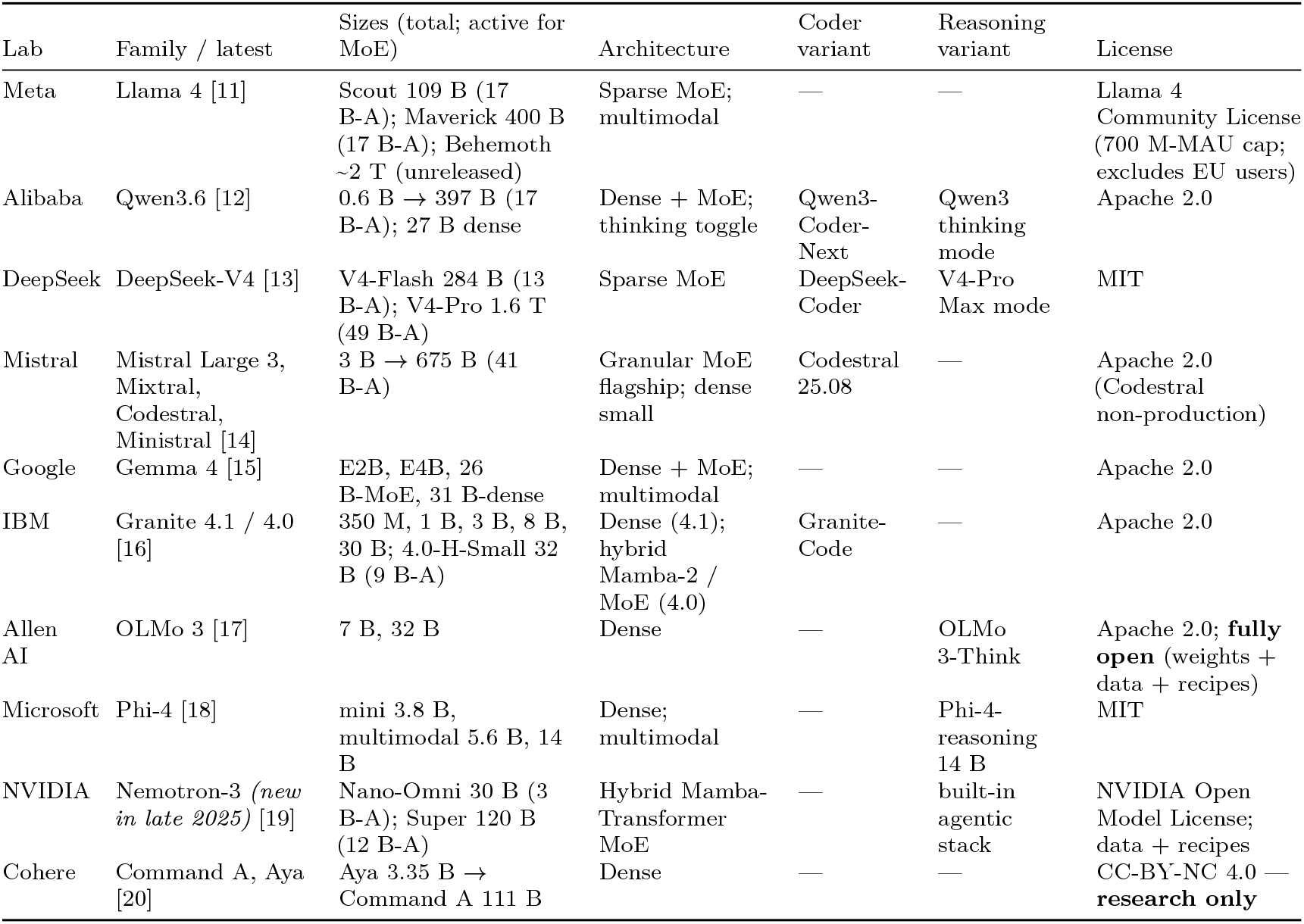
Major open-weight model families as of May 2026. Modern language models read input one piece at a time — each piece, called a *token*, is a short word or word-fragment — and predict the next piece by passing each token through a set of trainable mathematical operations. The **Architecture** column names the network structure: *Dense* — every internal weight is used on every token (the original transformer design, simple to deploy but compute per token scales with the model’s full size); *Sparse Mixture-of-Experts (MoE)* — the model is split into many parallel “expert” sub-networks plus a small “router” that, for each token, picks one or two experts while the rest stay idle (lets the model be very large in *total* parameters while spending the per-token compute of a much smaller model; every flagship released since late 2025 uses this design); *Hybrid Mamba-Transformer* — replaces some standard layers with *state-space* (Mamba) layers, so compute on a long input grows linearly with input length rather than quadratically (relevant now that flagships accept inputs of 500,000 to 1,000,000 tokens, a small book); *Multimodal* — accepts images, and sometimes audio, alongside text. **Sizes** reports parameters — the trained numerical weights inside a model, loosely analogous to synapse strengths — in billions (B) or trillions (T). *Total* parameters set disk and GPU-memory cost (a 70 B-parameter model occupies ∼35–40 GB at the standard 4-bit compression used for inference, or ∼140 GB at 16-bit “full precision”); for MoE models the column also reports *active* parameters per token (notation 17 B-A = 17 billion active per token), which set per-token compute cost — equal to total for dense models, much smaller than total for MoE because most experts are not triggered on a given token. **Coder variant** is a general model further trained on a large corpus of source code; coder variants beat their general siblings on bash, Python, and refactor tasks but lose on chat, math, and translation. **Reasoning variant** is a model that, before answering, writes out a step-by-step “chain of thought” that the user typically does not see; training rewards the model when its final answer is verifiably correct. Pioneered by DeepSeek-R1 (January 2025) and since copied across labs; reasoning variants score higher on math, code, and multi-step problems but take roughly 5×–50× longer per call (and cost 5×–50× more).

Several models are omitted from Table 1: Apple’s 3-billion-parameter on-device model ships only inside iOS and macOS 26, and xAI has openly released only Grok-1 (March 2024) and Grok-2.5 (August 2025) — everything newer is closed. Every flagship since late 2025 is a sparse MoE (see Table 1 legend), but dense persists below ∼40 B because it is simpler to deploy.

One practical reason narrows the choice for most biomedical labs – hardware required: the four size tiers in Table 1 map to four very different machines, from a $400–$600 consumer card for the smallest models to a multi-GPU server costing several hundred thousand dollars for the trillion-parameter tier (Table 2).

**Table 2.** GPU options and ballpark May 2026 US street prices for running each Table 1 model class locally at 4-bit compression. Where multiple cards are needed, the listed price is the per-card cost.

| Model class (Table 1 example) | GPU memory needed | GPU options (May 2026 USD) | Refs |
| --- | --- | --- | --- |
| **7–8 B dense** (Phi-4, Granite 8B, OLMo 3 7B) | ~5–6 GB | NVIDIA RTX 5060 Ti 16 GB (~\$560); RTX 4060 Ti 16 GB (~\$430); RTX 5070 12 GB (~\$635); Apple M4 Mac mini 16 GB unified (~\$600) | [21,22] |
| **27–32 B dense** (Qwen3.6-27B, Gemma 4 31B, OLMo 3 32B) | ~17–22 GB | NVIDIA RTX 5090 32 GB (~\$2,900–\$3,500, street price 50–75 % over \$1,999 MSRP); RTX 4090 24 GB (~\$1,500–\$2,200, EOL Oct 2024); RTX A5000 24 GB (~\$700–\$1,400 used); Apple Mac Studio M3 Ultra 96 GB unified (~\$4,000) | [21,23,22] |
| **100–400 B MoE** (Llama 4 Scout 109B, Maverick 400B) | ~55–250 GB | NVIDIA RTX Pro 6000 Blackwell 96 GB (~\$8,500); 2× RTX 5090 (~\$6,000–\$7,000 total); RTX 6000 Ada 48 GB (~\$6,800); H100 80 GB (~\$25K–\$33K); AMD Instinct MI300X 192 GB (~\$15K–\$20K, OEM-only); Apple Mac Studio M3 Ultra 256 GB unified (~\$9,500) | [21,23,24,25,22] |
| **Trillion-parameter MoE** (DeepSeek-V4-Pro 1.6T) | ~700+ GB | NVIDIA H200 141 GB (~\$31K–\$40K each); B200 192 GB (~\$35K–\$55K each); 8-GPU DGX H200 server (~\$350K–\$500K); GB200 NVL72 rack (~\$3M+); cloud rental on B200 ~\$2.25–\$16/GPU-hour | [24,26] |

Note: Spring 2026 prices are dominated by an ongoing memory shortage; consumer NVIDIA 50-series cards and high-RAM Apple Mac Studio configurations currently sell 30–75 % above launch MSRP.

Given this (rapidly evolving) landscape we decided to do the following experiment: take a common sequencing data processing workflow and ask open models running on hardware accessible to an average research lab to design and execute the analysis. In doing so we experimented with a range of possibilities ranging from allowing open models to figure out everything by themselves to guiding them using a very detailed plan produced by commercial frontier models. We further complicated these tasks by simulating a variety of errors that may occur during workflow execution.

## Materials and Methods

### Hardware

We assembled five computers spanning the lab-scale hardware tier (Table 3) — a salvaged workstation (assembled by combining components from two Dell Precision 5820 workstations), a gaming-GPU desktop, two recent Apple-silicon MacBooks, and the NVIDIA Jetson AGX Orin 64Gb – a “RaspberryPi”-like NVIDIA offering that costs under $2,000 and has a small footprint, making it a lab-ready tiny-but-powerful workstation.

**Table 3.** Test machines used in this study.

| Computer | Manufacturer | Year released | RAM | OS | GPU |
| --- | --- | --- | --- | --- | --- |
| NVIDIA Jetson AGX Orin Developer Kit | NVIDIA | 2022 | 64 GB LPDDR5 (unified with GPU) | Ubuntu 22.04 LTS (NVIDIA JetPack 6) | Integrated Ampere, 2,048 CUDA cores + 64 Tensor cores |
| RTX 5080 desktop | Dell | 2025 (GPU) | 128 GB DDR5 (system) | Linux (Ubuntu) | NVIDIA RTX 5080, 10,752 CUDA cores, 16 GB GDDR7 |
| MacBook Pro M4 Pro (48 GB) | Apple | 2024 | 48 GB LPDDR5X (unified with GPU) | macOS Sequoia 15.6 | Apple M4 Pro integrated GPU, 16 to 20 cores |
| MacBook Air M4 (24 GB) | Apple | 2025 | 24 GB LPDDR5X (unified with GPU) | macOS Sequoia 15.6 | Apple M4 integrated GPU, 10 cores |
| 2× NVIDIA RTX A5000 desktop | Dell | 2021 (GPUs) | 256 GB DDR4 (system) | Linux (Ubuntu 25.10) | 2× NVIDIA RTX A5000, 8,192 CUDA cores each, 24 GB GDDR6 each (48 GB total) |

System RAM listed for the two Linux desktops reflects the build configuration; for inference workloads the relevant memory is the GPU VRAM (last column). For the Jetson and the MacBook, RAM is unified between CPU and GPU and the model can use up to roughly the listed RAM minus the operating-system reservation.

A model’s GPU-memory footprint scales with its parameter count: at the 4-bit quantization used for all local runs here, roughly 0.5 GB per billion parameters. The plan-granularity-gradient sweep was executed on the **RTX 5080 desktop** with six 2026-release open-weight implementers — `gemma4:26b`, `gemma4:e4b`, `glm-4.7-flash`, `gpt-oss:20b`, `qwen3.6:27b`, and `qwen3.6:35b-a3b` — chosen so the family covers small dense (4 B effective), mid-size dense (26–27 B), and small-active sparse-MoE architectures within or just above the 5080’s 16 GB VRAM ceiling at 4-bit. The strongest implementer (`qwen3.6:27b`, 17 GB at 4-bit) is then cross-platform validated on the Jetson AGX Orin (64 GB unified memory), MacBook Pro M4 Pro (48 GB unified memory), MacBook Air M4 (24 GB unified memory — tight, sits within ∼1 GB of the macOS reservation), and 2 × RTX A5000 (48 GB total VRAM); the first three host the model comfortably while the Air spills to swap when background processes claim extra memory. The Anthropic API models (`claude-opus-4-7` as plan author throughout, plus `claude-sonnet-4-6` and `claude-haiku-4-5` as confirmation implementers) run remotely and are independent of host GPU.

### Data

We selected a small dataset [27] derived from our previous work on the analysis of mutational patterns in human mitochondria [28]. It contains four deeply downsized paired-end Illumina samples derived from blood and cheek tissues of a mother–child pair. These reads carry two fixed changes and a low-frequency variant in the child’s cheek sample — a heteroplasmy example.

### Workflow

We developed a simplified version of a haploid variant-calling workflow (original: [29]) that omits pre-processing steps before variant calling and the variant-annotation phase [30,31]. The structure of the workflow is shown in Fig. 1 below.

**Figure 1:**
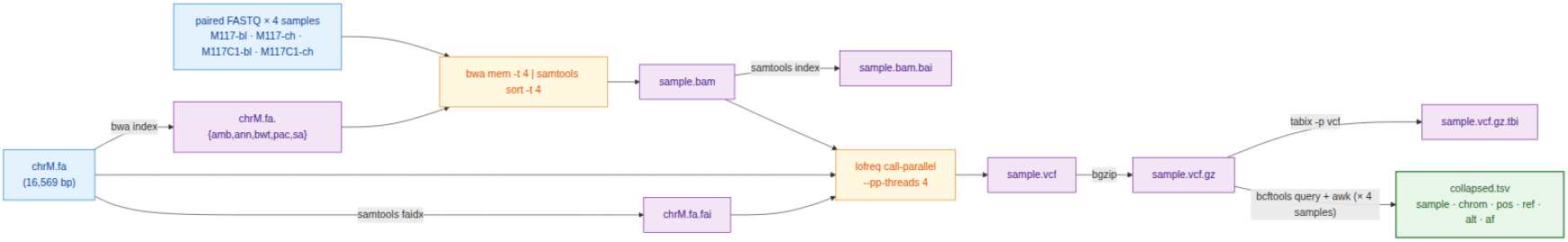
Data flow of the simplified mtDNA variant-calling workflow used in this study. Per-sample steps (alignment through tabix) run independently for each of the four samples; the only inter-sample dependency is the final bcftools query + awk fan-in that builds collapsed.tsv.

### Plans

We asked the free implementers to perform variant calling under conditions of varying recipe detail — from no plan at all to a very-detailed step-by-step specification (Table 4; full text representation of each plan is in the Supplement). Every plan was authored by claude-opus-4-7 in the planner role of the recipe-implementer split. Briefly, Opus wrote v1 (the lean reference recipe); v2 (a hyper-detailed recipe authored in response to v1’s poor implementer performance — see Results); v1.25 and v1.5 (controls that localize which part of v2 carries the score improvement); v1g (a robustness check authored mechanically from the Galaxy IUC tool registry rather than by Opus); v0.5 (a no-recipe condition with only the tool order); and v2_defensive (v2 plus an error-handling skeleton).

**Table 4.** Plan variants used in this study. Plans of increasing granularity, plus two “no-plan” controls. Sizes are bytes; the verbatim text of every plan is reproduced in the Supplement.

| Plan | Size | Short summary | Hypothesis tested |
| --- | --- | --- | --- |
| **Track B** | 0 | Problem statement + tool inventory only; no implementation hints | How much can a model do from scratch? |
| **v0.5** | 1.4 KB | Track B + one line giving the tool order: `bwa` → `samtools` → `lofreq` → `bcftools` → `awk` | Does sequencing alone help local models? |
| **v1 (lean)** | 3.1 KB | Numbered bullets naming tools and key flags; no exact command lines | Reference lean plan |
| **v1.25** | 3.1 KB | v1 + the exact `lofreq call-parallel` command line | Does one full command bridge the v1 → v2 gap? |
| **v1.5** | 1.3 KB | v2 with every prose paragraph and Gotchas block stripped — code fences only | Are the prose explanations load-bearing or decorative? |
| **v1g** | 4.2 KB | v1 + a Galaxy-IUC-mechanical `lofreq` snippet (extracted from the IUC tool registry [32]) | Can a tool registry replace a human plan author? |
| **v2 (detailed)** | 4.6 KB | Every step gives the exact command line plus Gotchas block | Reference detailed plan |
| **v2_defensive** | 6.5 KB | v2 + `try()` helper, output validation after every step, retry-once, per-sample isolation, structured failure log | Does explicit error-handling prose make implementer scripts defensive against runtime tool failures? |

Each model is run under one of two conditions, which we call tracks. In Track A, the model receives a written plan (any row in Table 4 except *Track B*) together with the tool inventory and writes `run.sh` from the plan. In Track B, the model receives the problem statement and the tool inventory only — no plan — and writes `run.sh` from scratch. Track B exists to bound how much of the implementer’s behavior comes from the model’s prior training versus from the supplied recipe; every other plan in Table 4 is a Track A condition.

### Error simulation

To test how each model handles tool failures, we wrapped bwa and lofreq in short shell scripts (“shims”) that imitate the real binaries on disk. Before each run, the harness writes the two shims into a per-run directory and adds that directory to the front of `PATH` — the colon-separated list of directories the shell consults when it needs to find an executable. When the script types `bwa mem`, the shell finds the shim first and runs it instead of the real bwa. Each shim reads two environment variables: `EVAL_INJECT_PATTERN` names the failure type and `EVAL_INJECT_TARGET` names which of bwa or lofreq the failure applies to. For every untargeted invocation the shim passes the call straight through to the real binary; for the targeted invocation it injects the failure and then either passes through or stops. The model’s `run.sh` does not know it is being intercepted.

Seven failure patterns cover the kinds of breakage a real lab pipeline encounters (Table 5). Each pattern is paired with one tool (or both), so a full error-matrix cell is one (model × plan × pattern × target tool × seed) combination. We use seeds 42, 43 and 44, and run every cell on both the `v2` and the `v2_defensive` plan to measure how the defensive prose changes outcomes.

**Table 5.** Failure patterns injected by the PATH shims into bwa and / or lofreq.

| Pattern | Affected tool | What the shim does |
| --- | --- | --- |
| `flake_first_call` | both | First call exits 1; later calls pass through. |
| `one_sample_fails` | both | Shim exits 1 only when `M117C1-ch` appears in the command line; the other three samples run clean. |
| `slow_tool` | both | Sleep 30 s, then run the real tool — output identical, just delayed. |
| `stderr_warning_storm` | both | Dump 200 lines of `WARNING:` text to stderr, then run the real tool — output is correct but the stderr looks alarming. |
| `missing_lib_error` | both | Exit 127 without invoking the real tool, mimicking a missing shared library (`error while loading shared libraries: libhts.so.3: cannot open shared object file`). |
| `silent_truncation` | lofreq only | Run the real lofreq, then truncate the `-o` output file to zero bytes — lofreq returns 0 but the VCF is empty. |
| `wrong_format_output` | lofreq only | Run the real lofreq, then strip every variant line from the VCF; only the header survives — the file still parses, but contains no calls. |

Beyond the variant-overlap score (defined in *Scoring* below), we score each error-matrix cell on three error-handling metrics — referred to here as the **handle category, recover**, and **diagnose** (fields `m_handle`, `m_recover`, `m_diagnose` in the per-cell score JSON). The handle category classifies the script’s overall response as `crash` (exited non-zero with no failure log), `propagate` (the failure passed through to a downstream step), `partial` (the script aborted but recorded the failure structurally), or `recover` (the script completed successfully despite the injection). The recover metric is binary: did the script produce the count of valid VCFs that the failure pattern allows? For `one_sample_fails` the best achievable is 3 (the bad sample’s VCF is correctly absent); for `silent_truncation`, `wrong_format_output` and `missing_lib_error` the best is 0 (the script should detect the broken or missing output and skip it); for the remaining three patterns the best is 4. The diagnose metric is binary: did the script announce the failure — through a populated `failures.log`, a summary line on stderr, or a sample-name-and-failure-word pair in its output — or did it stay silent? All three metrics are defined in `score/score_run.py:error_handling`.

### Scoring

We score every run on a single primary metric — the **variant-overlap (Jaccard) score** — computed automatically by `score/score_run.py` from the script’s outputs and the canonical answer key. The variant-overlap score is the per-sample Jaccard overlap of `(chrom, pos, ref, alt)` between the script’s PASS-or-unfiltered records and the answer key, requiring matched allele frequencies within ±0.02 and averaged over the four samples; it ranges from 0 (no overlap) to 1 (every call correct). We gate the score on script execution: a run whose `bash run.sh` invocation does not return exit 0 within the 600-second wall-clock budget contributes its produced VCFs as-is to the average, and any missing VCF counts as zero overlap on its sample. A model that uses a different but valid tool from the one named in the plan — `bcftools mpileup` in place of lofreq, for example — gets credit for the variants it called, not for the tool it chose.

For the error-injection cells (described under *Error simulation*) we add the three error-handling metrics defined above (handle category, recover, diagnose). The per-run `score.json` also records additional bookkeeping fields — token counts, USD cost (Anthropic only — Ollama is free), generation and execution times, file-schema completeness, and shellcheck/idempotency status — that are written for reproducibility but are not analyzed in the Results below.

### Code, data, and artifact availability

Every step of this study — plan authoring, harness construction, sweep execution, scoring, figure generation, and manuscript drafting — was driven from Anthropic’s **Claude Code** command-line agent (https://claude.com/claude-code). The manuscript was then extensively modified, clarified, and otherwise edited by the author. Plans v1 through v2_defensive were authored by claude-opus-4-7 inside Claude Code; the harness (`harness/run_one.py`, `harness/error_matrix.py`, `harness/sweep_local.py`) and the scoring code (`score/score_run.py`, `score/aggregate.py`) were written by claude-opus-4-7 in the same session; the multi-platform sweeps (RTX 5080, NVIDIA Jetson AGX Orin, MacBook Pro M4 Pro, MacBook Air M4, 2 × RTX A5000) were dispatched as Claude Code sessions on each machine, with results pushed back as pull requests; the figures in this manuscript were generated by `scripts/make_figures.py` (Altair / Vega-Lite).

All artifacts — the seven plans, the prompt templates, the four mtDNA samples and the chrM reference, the canonical workflow and reference VCFs, the harness with the PATH-shim error-injection scaffolding, the per-run scorer, the aggregate score table, the per-cell error-matrix JSONLs from every platform, the figure-generation scripts, the manuscript source, and the rendered PDF — are released under MIT at https://github.com/nekrut/LLM-eval-paper. The repository’s `README.md` documents how to re-run the experiment on any new model in three commands once Ollama is installed; `setup/RUN_ON_MACOS.md` covers the Apple-silicon-specific path including a decision tree for the 24 GB MacBook Air’s tight-memory regime.

## Results

### All you need is a plan

Every (model × plan) condition reported below was run three times with different random seeds (42, 43, 44). A seed is the integer that controls the model’s token-sampling stochasticity — the same prompt-and-seed pair produces reproducible output, while different seeds produce independent samples from the same underlying probability distribution, letting us distinguish robust success (all three seeds pass) from intermittent success (only one or two seeds pass).

Five of six 2026-release open-weight implementers reach mean score 1.000 on the v2 plan, the most detailed Opus-authored recipe. We ran the six implementers on the RTX 5080 against every plan in Table 4 — Track A (with plan) and Track B (no plan) — with three seeds per (model × plan) condition, scoring the variant-overlap as the share of called variants matching the truth set within a 0.02 allele-frequency window (Methods, *Scoring*). The five passing implementers are qwen3.6:27b, qwen3.6:35b-a3b, gemma4:26b, gemma4:e4b, and glm-4.7-flash; the sixth, gpt-oss:20b, reaches mean score 0.67. The recipe-implementer split is operational on commodity hardware: a frontier-tier author writes the recipe once and a 17 GB open-weight implementer reproduces frontier accuracy at zero per-call cost on a desktop GPU.

### Plan granularity has high impact

To localize what an implementer needs from the recipe, we constructed plans of increasing detail (Fig. 2). At the lean end, claude-opus-4-7 first wrote v1: a numbered bullet list naming each tool and its key flags but no exact command lines (see Supplement). v1 produces an all-or-none response — only qwen3.6:27b reaches score 1.000 reliably across all three seeds; glm-4.7-flash passes two of three seeds (mean 0.667); the four other implementers fail at score ≈ 0. The detailed v2 recipe, written in response, recovers all six implementers (subject to the gpt-oss:20b budget effect noted above). To isolate which part of v2 is doing the work, we authored two intermediate plans: **v1.25** = v1 plus the single literal `lofreq call-parallel --pp-threads 4 -f data/ref/chrM.fa -o results/{s}.vcf results/{s}.bam` invocation, and **v1.5** = v2 with every prose paragraph and *Gotchas* block stripped, code fences only. Both v1.25 and v1.5 reproduce the v2 jump on the same dense ≥ 26 B set — qwen3.6:27b, qwen3.6:35b-a3b, and gemma4:26b reach 1.000 on both; the prose between the v1.25 line and the v2 prose is decorative for these implementers. The active ingredient is the literal command syntax. Track B (no plan) and v0.5 (no plan plus the tool order bwa → samtools → lofreq → bcftools → awk) confirm the floor: every open-weight implementer’s scores at 0 in both no-recipe conditions; supplying tool order without flags or arguments does not lower the threshold.

**Figure 2:**
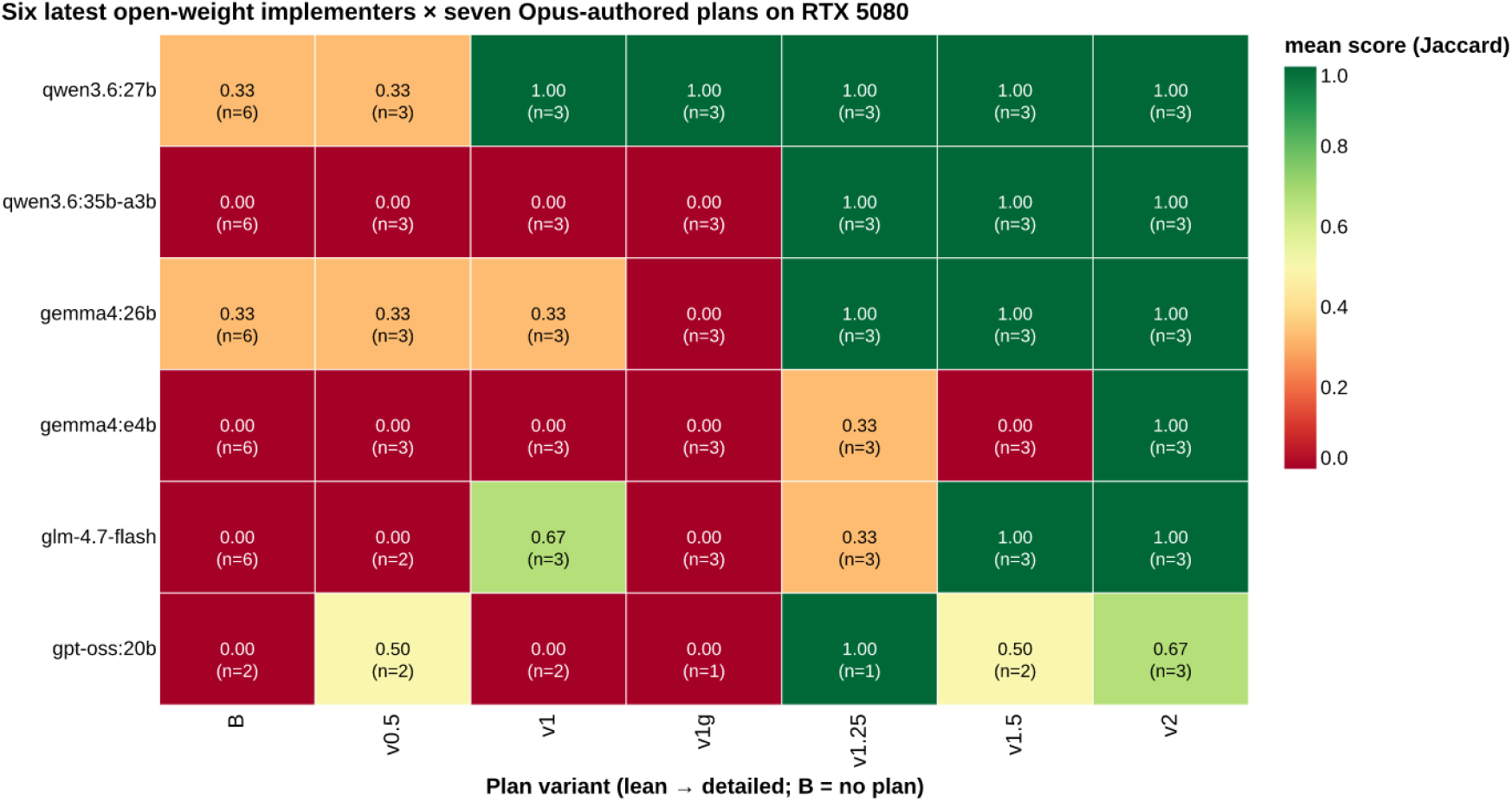
Mean score across three seeds for six open-weight implementers × seven Opus-authored plan variants on the RTX 5080. Columns are ordered by plan detail (left = no plan, right = most detailed). Anthropic API plan-author rows are not shown here because Anthropic models execute on Anthropic’s own hardware; they appear in the cross-platform error-injection panel below.

We tested an alternative source for the canonical command line with **v1g** = v1 plus the lofreq call-parallel snippet extracted mechanically from the Galaxy IUC tool registry [32] at commit `39e7456`, a hand-curated public corpus of bioinformatics tool wrappers. v1g does not reproduce the v1.25 lift — only qwen3.6:27b passes (3/3 seeds at 1.000); the other five implementers stay at score ≈ 0, indistinguishable from v1. The IUC snippet carries the same lofreq invocation but embeds it inside Galaxy’s macro infrastructure; that surrounding context appears to interfere with implementers other than qwen3.6:27b. The active ingredient is therefore the canonical Opus-authored single-line invocation as it appears in v1.25 and v2, not any presentation of the same command.

### qwen3.6:27b is the winner

Within the six-implementer lineup, qwen3.6:27b (Apache 2.0, Alibaba; 17 GB at 4-bit) is the only one that reaches score 1.000 on every v1 seed. The next-strongest, glm-4.7-flash (Z.ai; comparable size), passes two of three seeds for a mean of 0.667 — close enough to confirm the threshold response is not a single-model artifact, but not consistent enough to recommend the model for production. qwen3.6:27b also has a permissive Apache-2.0 license with no field-of-use restrictions, ships as a single-file `ollama pull qwen3.6:27b` deployment, and fits the 24 GB VRAM tier on Table 2 with room to spare. We therefore selected qwen3.6:27b as the protagonist for the cross-platform validation that follows.

### “Typical” lab hardware is sufficient

Hardware does not limit accuracy on the v2 plan — only execution (wall) time. We tested qwen3.6:27b on the four remaining platforms — Jetson AGX Orin, MacBook Pro M4 Pro, MacBook Air M4 (24 GB), and 2 × RTX A5000 — under the v2 plan; mean score 1.000 on every platform on every seed. However, wall-clock generation time differs sharply, and on the most memory-constrained platforms swap behavior dominates the variance (Table 6). The 2 × A5000 returns a generation in under half a minute (median 29 s), the M4 Pro and Jetson take 1.5 to 2 minutes (92 s and 105 s), and the RTX 5080 and MacBook Air sit between 5 and 7 minutes (302 s and 518 s, respectively); both reflect the model partially spilling out of dedicated VRAM (5080, 16 GB) or constrained unified memory (Air, 24 GB) into system RAM or swap, which Ollama handles automatically but slowly. The MacBook Air is the worst-case configuration in our lineup — qwen3.6:27b at ∼17 GB sits within ∼1 GB of the macOS reservation, so a single seed (43) where background processes consumed extra memory ran 16 × slower than its faster sibling (4,345 s vs 276 s on seed 42), and 47 × slower than the median MacBook Pro M4 Pro generation (92 s). The accuracy was unaffected; the wall time was not. None of the platforms is too slow to be useful for a working lab; pick the cheapest box that fits the model in dedicated or unified memory with headroom and the score is unchanged. For the tightest hardware tier (16–24 GB), `qwen3:14b` (∼9 GB at 4-bit) is a comfortable fallback — on the same MacBook Air it ran in median 99 s with no swap penalty (3/3 seeds, mean score 1.000).

**Table 6.** qwen3.6:27b on the v2 plan: median wall-clock time per generation (seconds), inter-quartile range or full range in brackets. **n** is the number of independent generations aggregated per platform — Jetson, MacBook Pro M4 Pro, and 2 × A5000 contribute n=36 from the 12 PATH-shim error-injection cells × 3 seeds (Methods, *Error simulation*); the RTX 5080 contributes n=6 from the v2 plan-gradient sweep with reruns; the MacBook Air M4 contributes n=3 from the quick-check sweep documented in setup/RUN_ON_MACOS.md. Score 1.000 on every cell across every platform.

| Platform | VRAM/UMA | Median wall time (s) | Range (s) | n |
| --- | --- | --- | --- | --- |
| 2× RTX A5000 | 48 GB total VRAM (fits) | 29 | [3–10] (IQR) | 36 |
| MacBook Pro M4 Pro | 48 GB unified (fits) | 92 | [91–136] (IQR) | 36 |
| NVIDIA Jetson AGX Orin | 64 GB unified (fits) | 105 | [98–107] (IQR) | 36 |
| RTX 5080 desktop | 16 GB VRAM (spills to RAM) | 302 | [288–322] (IQR) | 6 |
| MacBook Air M4 | 24 GB unified (tight; swap-sensitive) | 518 | [276–4,345] (full) | 3 |

### Small LLM matches accuracy of a frontier API model despite error-injection

qwen3.6:27b matches claude-opus-4-7 cell-for-cell on the error-injection matrix. To probe the error handling ability of qwen3.6:27b we ran the 36-cell error matrix (12 PATH-shim cells 3 × seeds; Methods, *Error simulation*) for both models on three platforms — Jetson, M4 Pro, and 2 × A5000 — under both the v2 baseline plan and the v2_defensive plan that adds a try() {…} helper, per-step output validation, and a structured failures.log (Methods, *Plans*). Each cell is classified into one of four “handle categories”: *crash* — the script exits non-zero with no failure log, leaving the lab user with a broken run and no diagnosis; *propagate* — the script keeps running past the failed step, feeding malformed or missing output to downstream steps and producing an unusable final result; *partial* — the script handles the failed step by skipping the bad sample, recording the failure in a log, and completing the rest; and *recover* — the script works around the failure (e.g. by retrying a problematic tool call) and produces output indistinguishable from an uninjected run. Both models behave identically at the cell level on every platform tested (Fig. 3, Table 7).

**Table 7.** Per-cell behavior signature and mean variant-overlap score for qwen3.6:27b and claude-opus-4-7 on the 36-cell error matrix (12 patterns × 3 seeds). Counts and mean score are identical across Jetson, MacBook Pro M4 Pro, and 2 × RTX A5000 for every (model, plan) row.

| Model | Plan | recover | partial | propagate | crash | mean score |
| --- | --- | --- | --- | --- | --- | --- |
| claude-opus-4-7 | v2 | 15 | 0 | 6 | 15 | 0.333 |
| claude-opus-4-7 | v2_defensive | 21 | 15 | 0 | 0 | 0.625 |
| qwen3.6:27b | v2 | 15 | 0 | 6 | 15 | 0.333 |
| qwen3.6:27b | v2_defensive | 21 | 15 | 0 | 0 | 0.625 |

**Figure 3:**
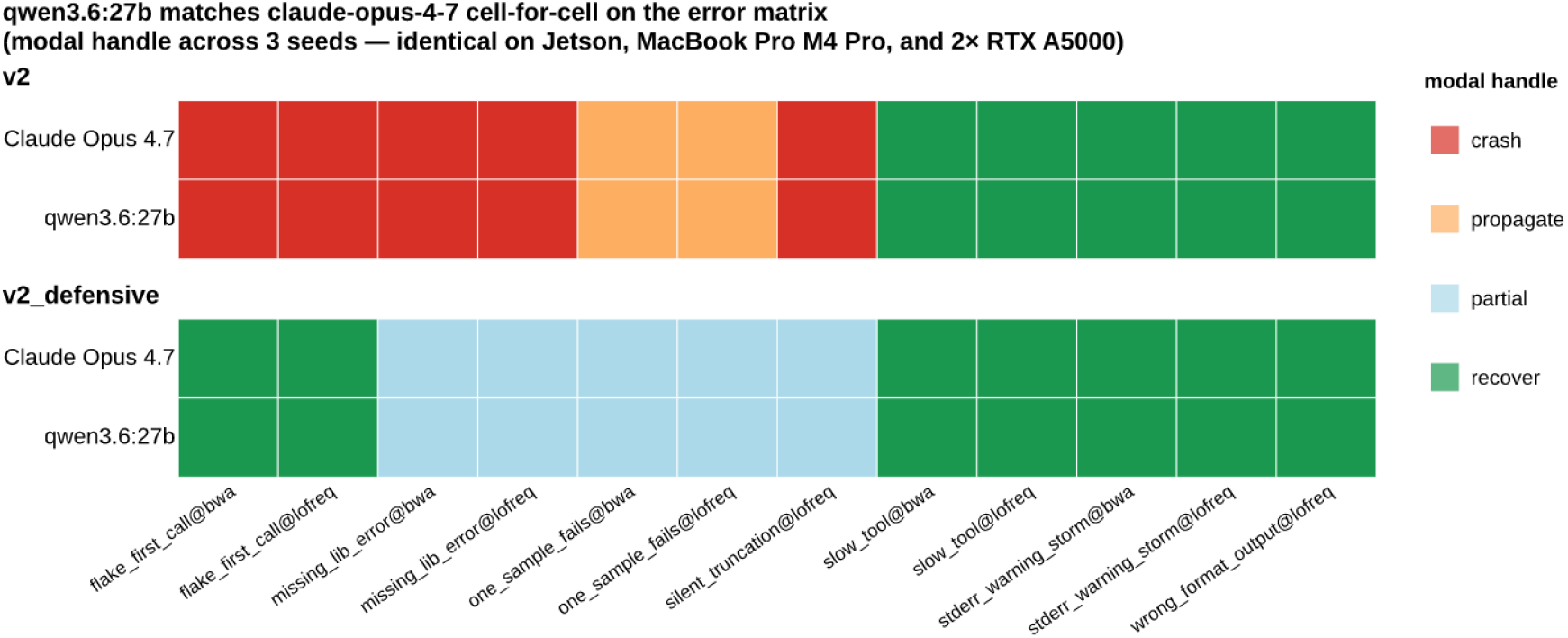
Per-cell modal handle category (recover / partial / propagate / crash) for claude-opus-4-7 and qwen3.6:27b across the 12-cell PATH-shim error matrix. Left panel = v2 baseline plan; right panel = v2_defensive plan. Rows within each panel are the two models. Cells are color-coded by the modal handle category across three seeds; the pattern shown is identical on the Jetson AGX Orin, MacBook Pro M4 Pro, and 2 × RTX A5000 platforms (verified for every one of the 48 cells), so a single cross-platform consensus is shown.

The error-injection result closes the loop on the recipe-implementer architecture: under both the happy-path plan and a defensive plan, a free, locally-runnable Apache-2.0 implementer handles the same workload as the frontier API author at the same accuracy.

### Cheap models offer an alternative

If local-model deployment is not an option, users can still cut per-call cost substantially by using cheaper commercial models as implementers — Opus authors the recipe once at premium rates, but Sonnet or Haiku then executes it many times at a fraction of Opus’s per-call price. Here we provide examples with claude-sonnet-4-6 and claude-haiku-4-5: run as implementers against the same 36-cell error matrix on Jetson and 2 × A5000, both produce the v2_defensive frontier signature of Opus and qwen3.6:27b — Sonnet at 21 recover / 14 partial / 0 propagate / 1 crash on Jetson and 21/15/0/0 on A5000 (mean score 0.625); Haiku at 21/15/0/0 on both platforms (mean score 0.625). Cross-platform timing for Anthropic models is not informative for hardware-throughput claims — the LLM call is remote — so we report these results as a commercial-fallback path for the implementer role, not as a hardware comparison.

## Discussion

A substantial fraction of routine biological data analysis will, within the next few years, run inside agentic tools — software environments where a large language model plans the work, calls external tools, executes code, reads the outputs, as well as iterates with minimal human intervention. Our group has been interested in the cost structure of this transition: who pays for the inference, and how often. The recipe-implementer split tested here is, in our view, the natural cost-control pattern for the transition.

A growing class of agentic tools — OpenCode (a terminal-based open-source agent from sst, supporting Anthropic, OpenAI, OpenRouter, as well as any local Ollama provider), Aider (a terminal coding agent with the same multi-provider model selection), Cline and its fork Roo Code (VS Code extensions), Continue (an IDE plugin for VS Code as well as JetBrains), Block’s Goose, Cursor, and Pi.dev, among others — share one design choice: the user picks the model. Most allow per-task or per-prompt model swapping, so the model that plans does not have to be the model that executes the next sub-step.

The recipe-implementer split maps onto these tools’ model-swap feature directly. Briefly, plan once with the most capable model the budget allows (claude-opus-4-7 in our experiments, but DeepSeek-V4-Pro Max or any equivalent frontier model would work the same way); then switch the agent to a local Ollama model — qwen3.6:27b at 17 GB, qwen3:14b at 9 GB for tighter memory tiers — for every implementation step that follows. The authoring cost is paid once per recipe (depending on the complexity of the workflow); the per-call inference cost drops to zero. For a lab that runs the same workflow on hundreds or thousands of samples a year, the difference is two orders of magnitude in cumulative LLM spend.

Two appliances stand out, in our view, as the recommended lab-bench computational hardware for the implementer side: the **NVIDIA Jetson AGX Orin Developer Kit** (under $2,000, ∼25 W power draw, 64 GB unified memory) and the **Apple Mac Mini** configured with maximum unified memory (M4 Pro at 64 GB, ∼$2,000). Both reproduce frontier accuracy on the v2 plan, both run qwen3.6:27b out of the box, as well as fit on the corner of a benchtop in a small-footprint, near-silent, low-power form factor that does not compete with wet-lab equipment for space or noise budget. The Jetson runs the standard Linux toolchain through NVIDIA’s JetPack image; the Mac Mini ships ready to use on macOS. Other tested configurations also work — a single $400–$600 consumer GPU (RTX 5060 Ti or 5070 with 16 GB VRAM, or a used A4000 with 16 GB VRAM; Table 2) runs qwen3.6:27b on the lean v1 plan, and a recent Apple-silicon MacBook with 48 GB or more of unified memory runs it on v2; even a 24 GB MacBook Air deploys the pipeline through the qwen3:14b fallback for the tight-memory tier. We tested all four configurations and found accuracy unchanged across them — only wall time varies. The implication for a working lab is direct: most labs already own something that works.

There are also several potential limitations of our approach. First, our study covers a single workflow — per-sample mtDNA variant calling on a 16.6 kb reference; the recipe-implementer split may behave differently on larger references, longer pipelines, or workflows depending on numerical algorithms beyond shell-stitched binaries (statistics, image processing, single-cell analyses). Second, claude-opus-4-7 is an API-only model and reproducibility of the recipes themselves depends on Anthropic’s continued availability of that model identifier; the recipes can be archived as text (they are short, human-readable, and we include all seven verbatim in the Supplement), however the act of authoring a new recipe for a new task remains tied to a closed API. Third, the PATH-shim error-injection harness probes failure modes that touch tool stdout/stderr, exit codes, as well as output-file shape; it does not probe failures depending on tool internals. The implementer-quality story is therefore strictly about recoverable failures detectable at the script level.

## Acknowledgements

This study was supported by NIH grants U24HG006620, U24AI183870, U24HG010263, OT2OD037936, and R01GM151683, as well as NSF award 2419522. The author thanks the Galaxy team for support and advice. The content is solely the responsibility of the author and does not necessarily represent the official views of the NIH or NSF.

## Supplement

### Plan files

The full text of each plan file referenced in Table 4. Each plan is reproduced verbatim from `plan/` in the repository and shown inside a fenced Markdown block to preserve its original heading hierarchy and code formatting.

### Plan v1 (lean) — plan/PLAN_v1.md

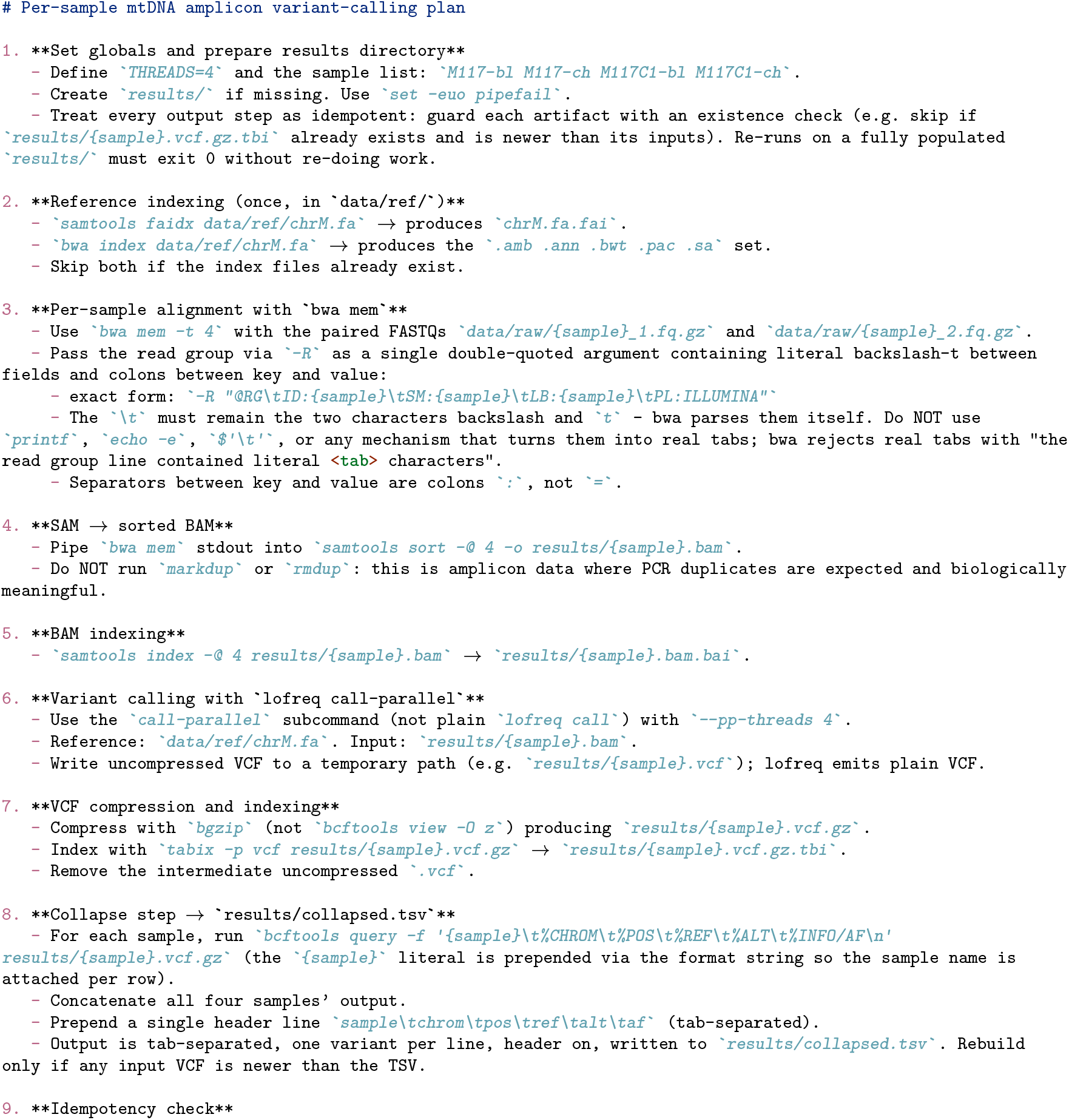
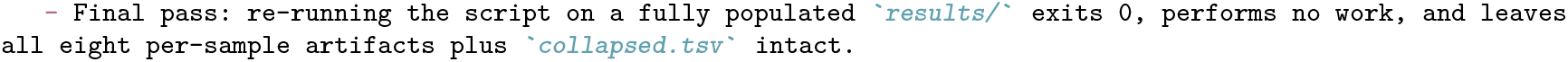

### Plan v1.25 — plan/PLAN_v1p25.md

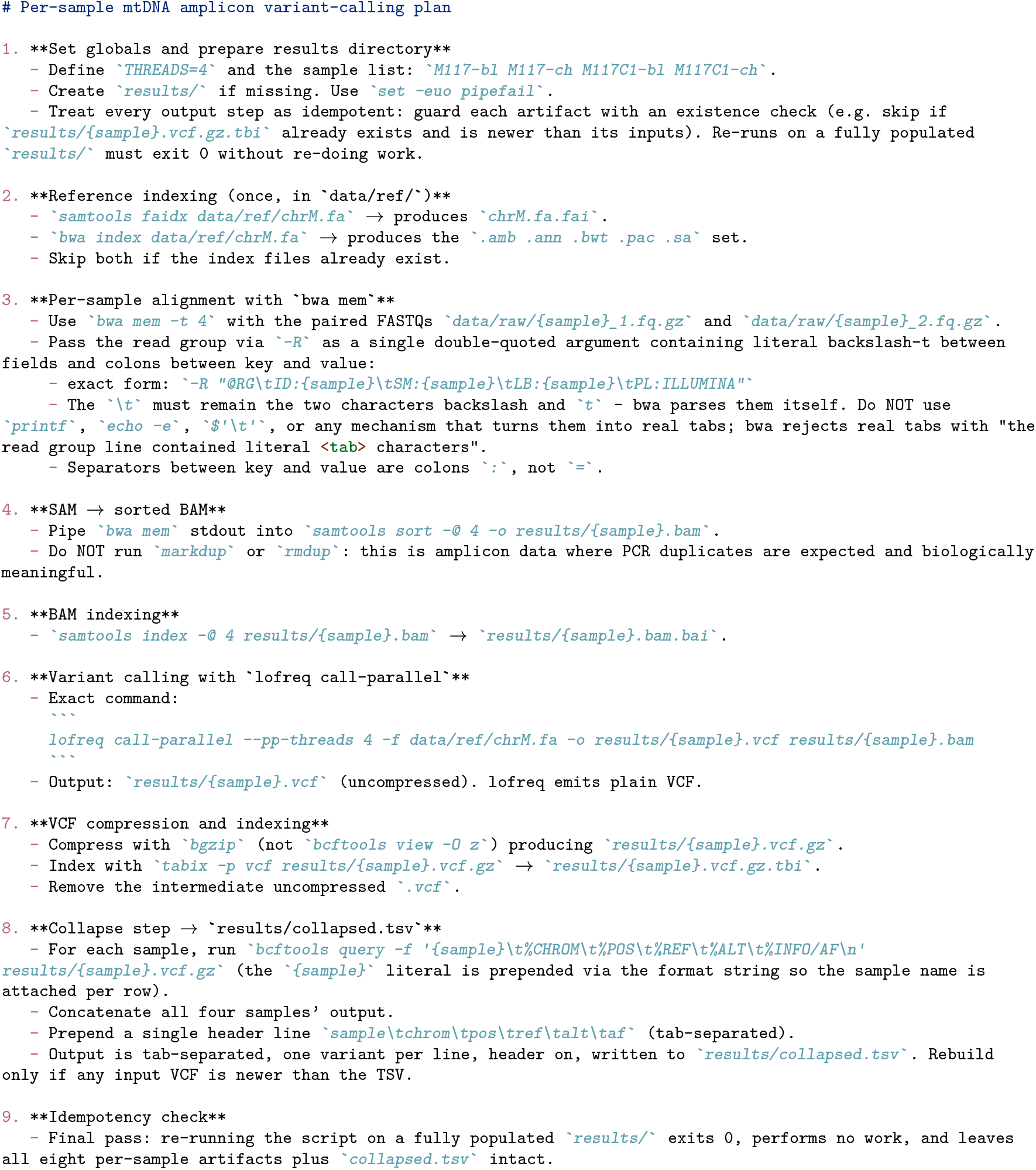

### Plan v1.5 — plan/PLAN_v1p5.md

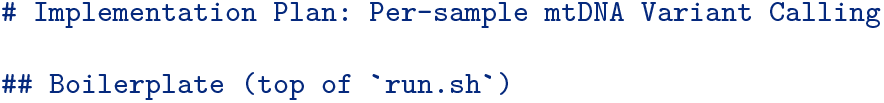
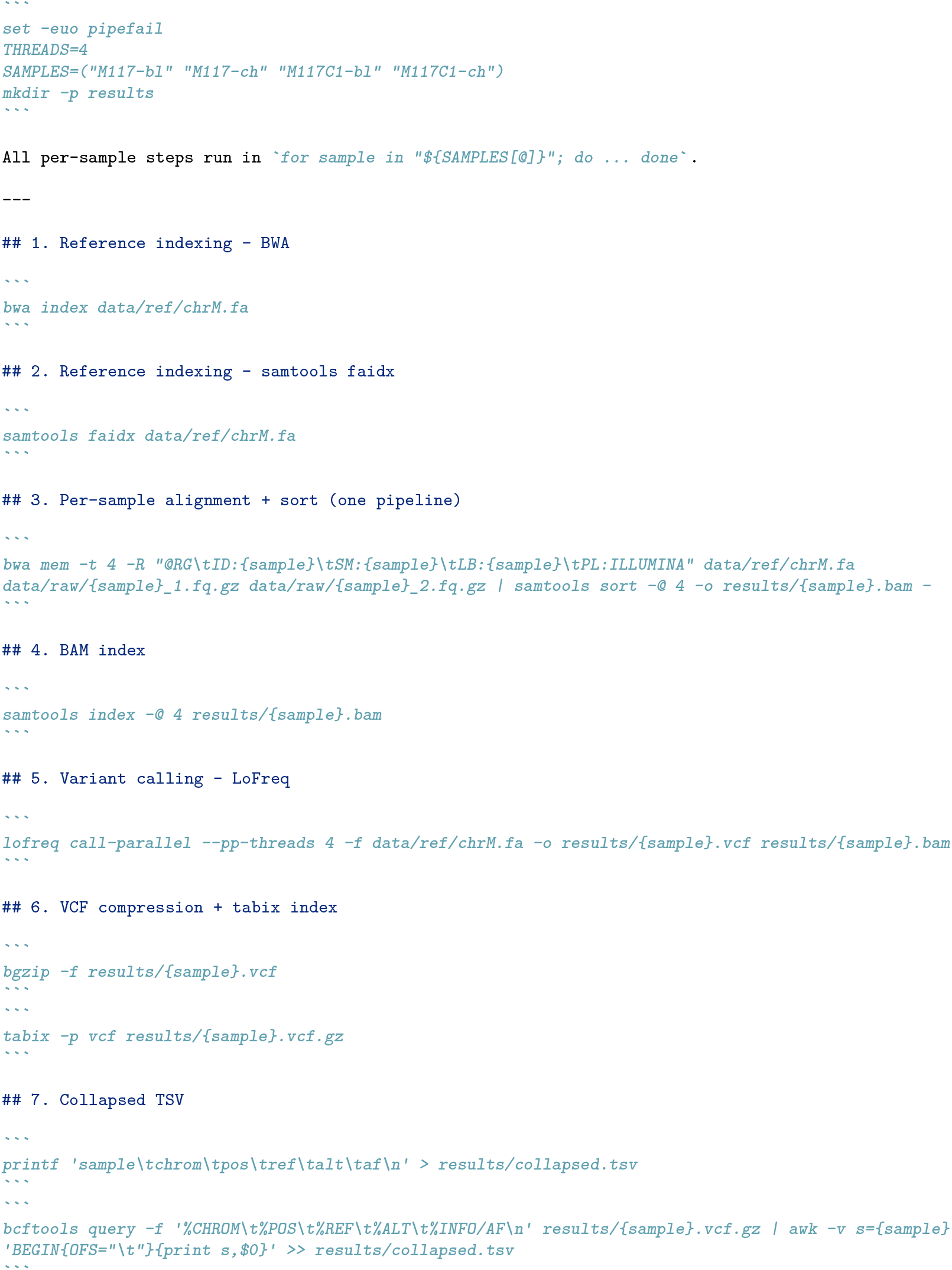

### Plan v1g — plan/PLAN_v1g.md

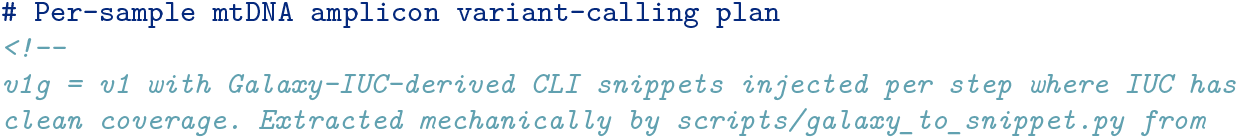
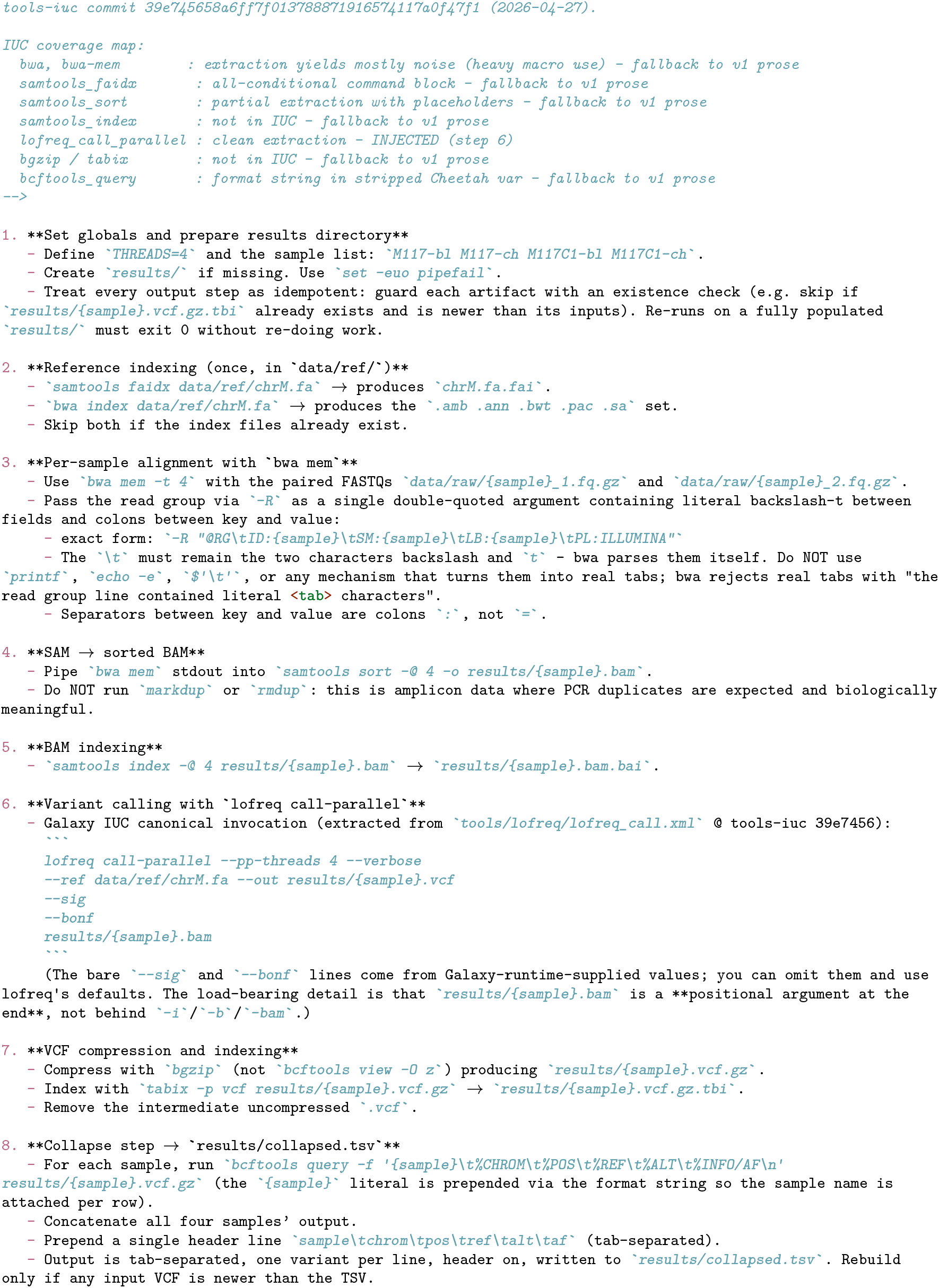
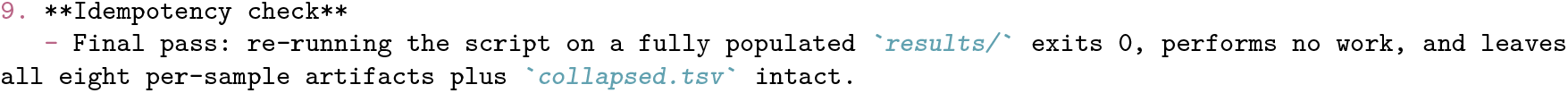

### Plan v2 (detailed) — plan/PLAN.md

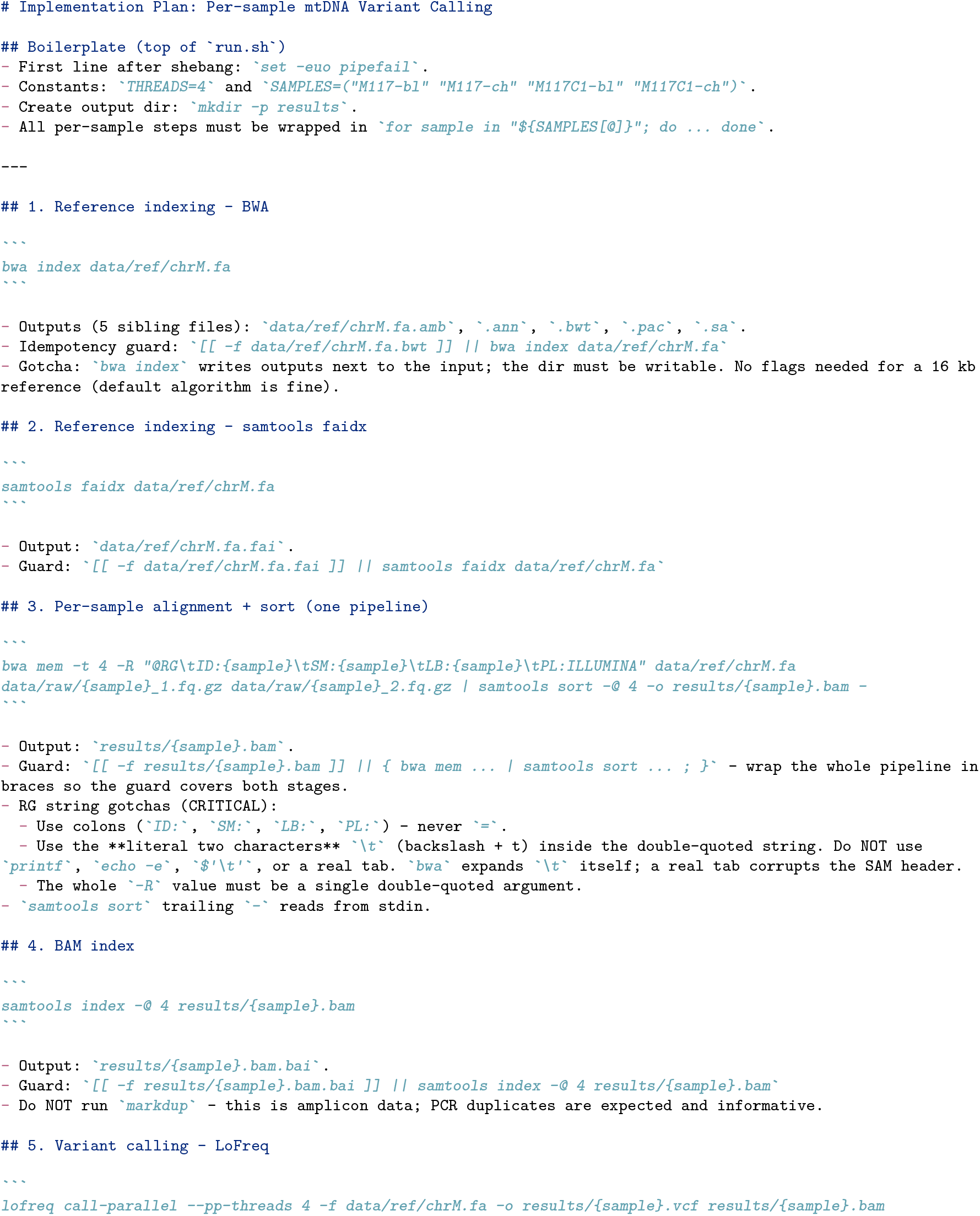
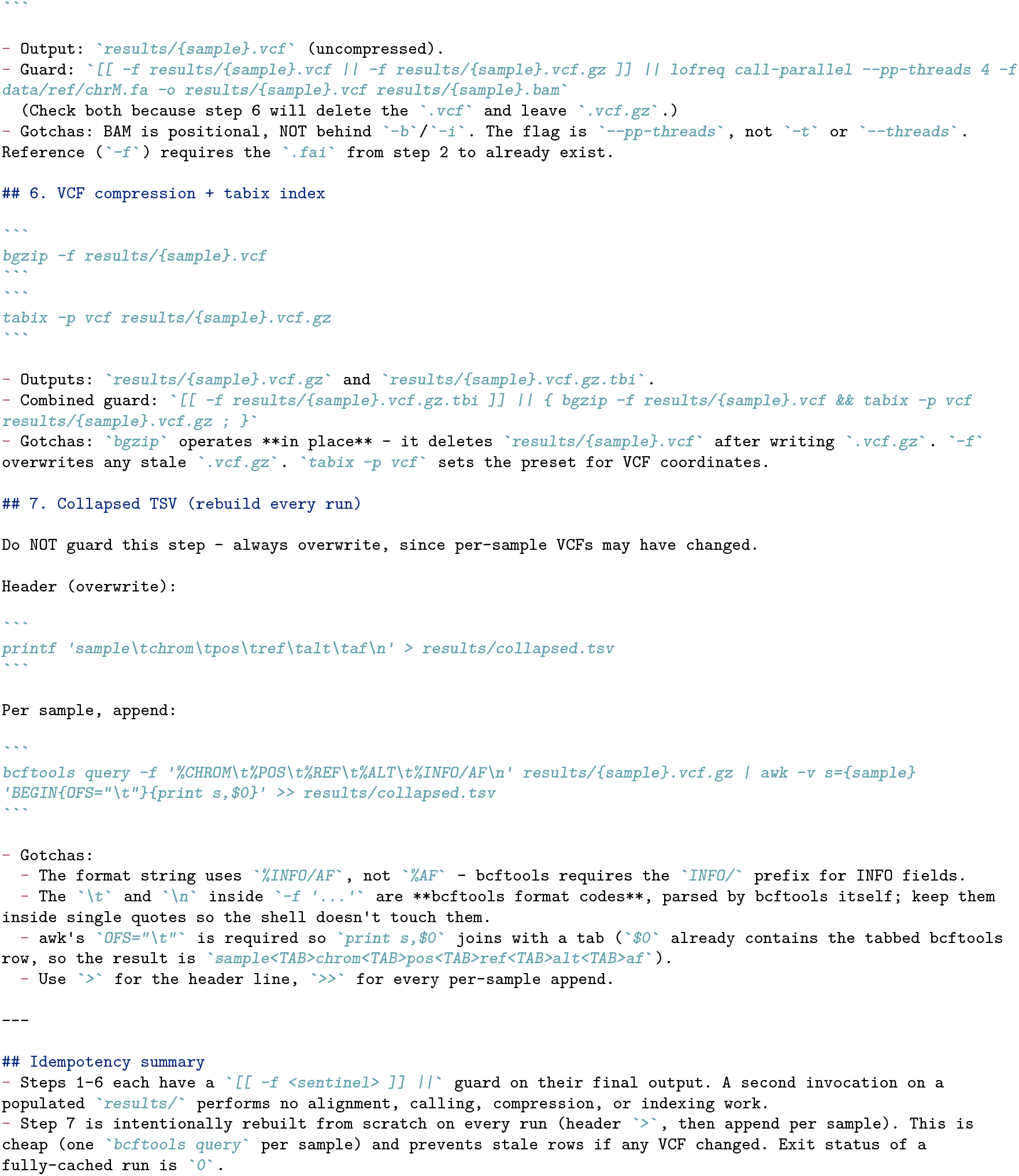

### Plan v2_defensive — plan/PLAN_v2_defensive.md

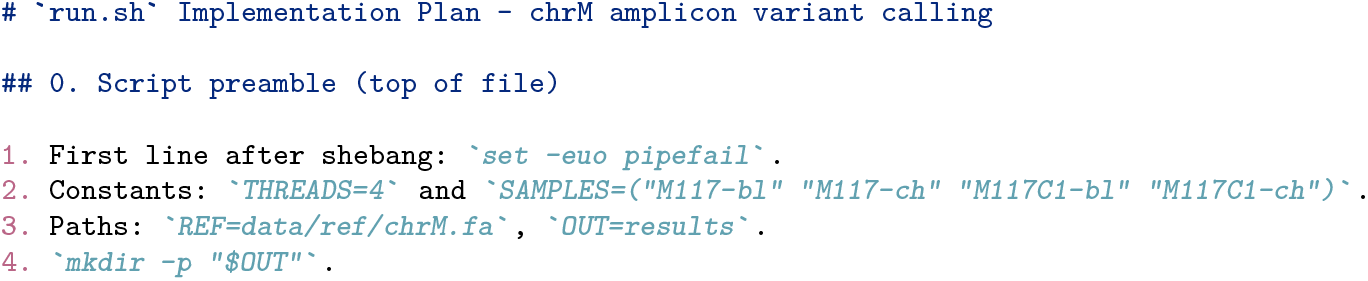
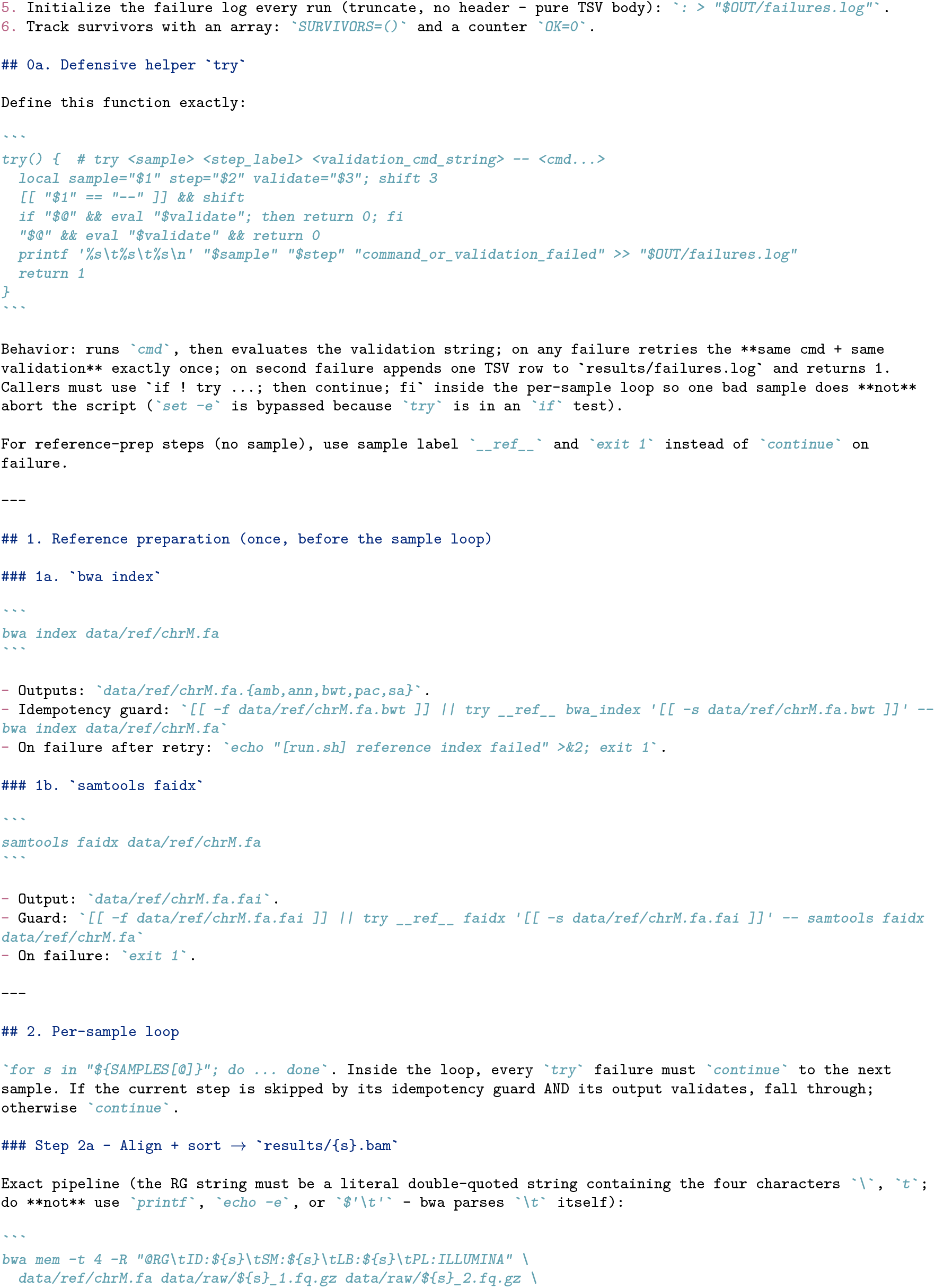
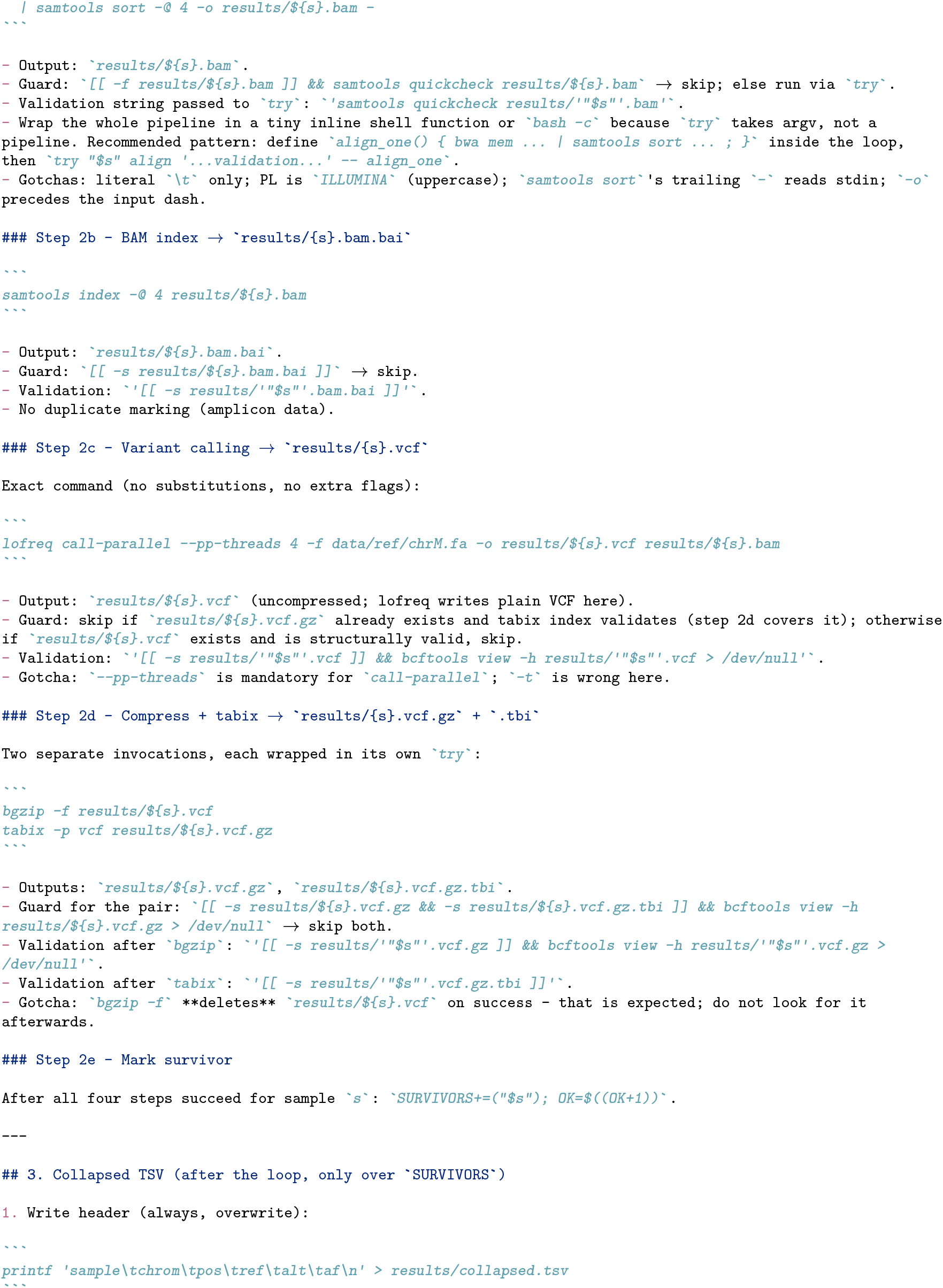
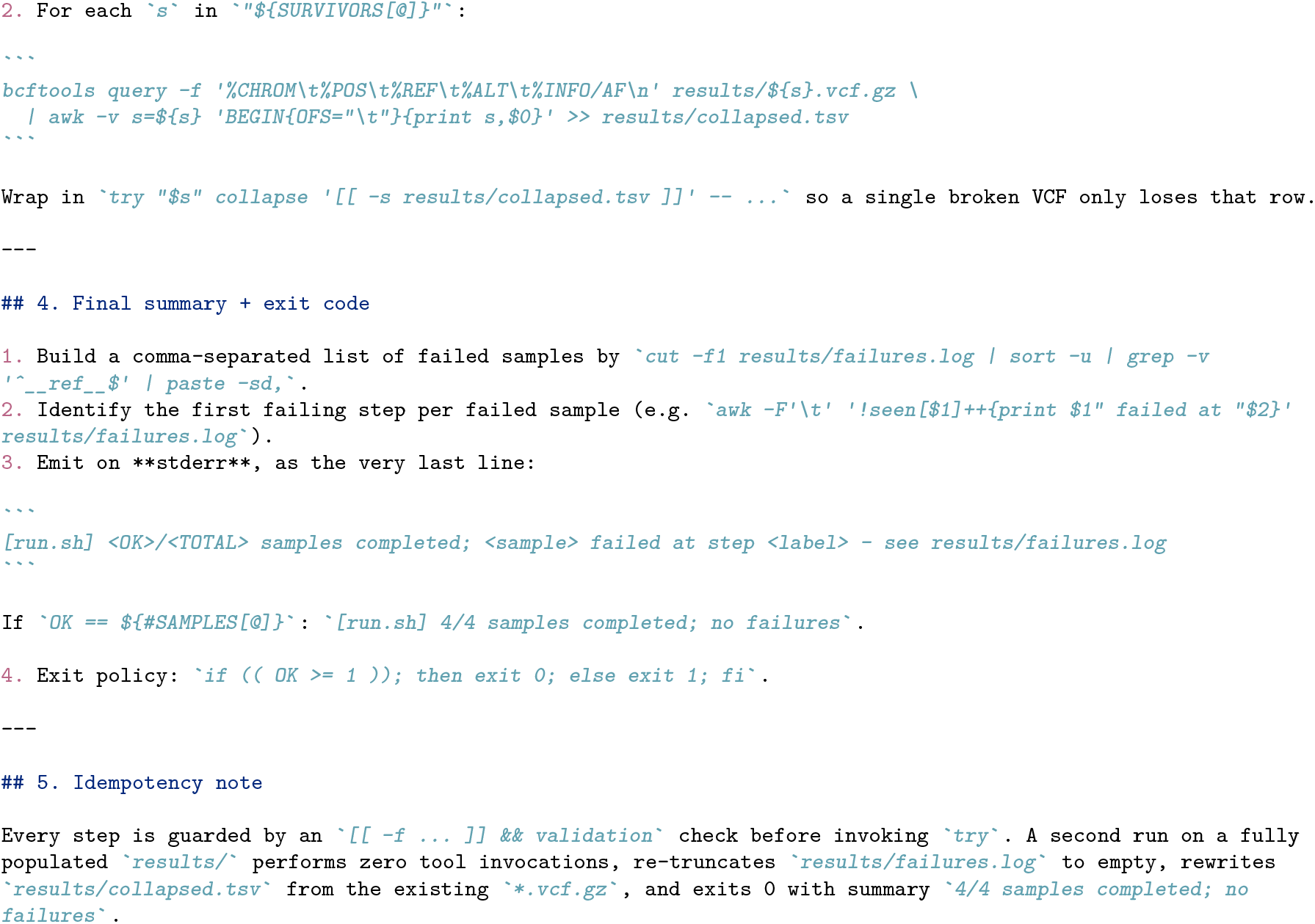

## References

[1] Jin Q, Yang Y, Chen Q, Lu Z. GeneGPT: augmenting large language models with domain tools for improved access to biomedical information. Bioinformatics. 2024;40(2):btae075. doi:10.1093/bioinformatics/btae075. PMID:38341654.

[2] Shang X, Liao X, Ji Z, Hou W. Benchmarking large language models for genomic knowledge with GeneTuring. Brief Bioinform. 2025;26(5):bbaf492. doi:10.1093/bib/bbaf492.

[3] Tang X, Qian B, Gao R, Chen J, Chen X, Gerstein MB. BioCoder: a benchmark for bioinformatics code generation with large language models. Bioinformatics. 2024;40(Suppl_1):i266–i276. doi:10.1093/bioinformatics/btae230. PMID:38940140.

[4] Sarwal V, Andreoletti G, Munteanu V, Suhodolschi A, Ciorba D, Bostan V, Dimian M, Eskin E, Wang W, Mangul S. BioLLMBench: a benchmark for large language models in bioinformatics. bioRxiv; 2023. doi:10.1101/2023.12.19.572483.

[5] Rajesh V, Siwo GH. Out-of-the-box bioinformatics capabilities of large language models (LLMs). bioRxiv; 2025. doi:10.1101/2025.08.22.671610. PMID:40909484.

[6] Mitchener L, Laurent JM, Andonian A, Tenmann B, Narayanan S, Wellawatte GP, White A, Sani L, Rodriques SG. BixBench: a comprehensive benchmark for LLM-based agents in computational biology. arXiv:2503.00096; 2025. https://arxiv.org/abs/2503.00096

[7] Su H, Long W, Zhang Y. BioMaster: multi-agent system for automated bioinformatics analysis workflow. bioRxiv; 2025. doi:10.1101/2025.01.23.634608.

[8] Mehandru N, Hall AK, Melnichenko O, Dubinina Y, Tsirulnikov D, Bamman D, Alaa A, Saponas S, Malladi VS. BioAgents: bridging the gap in bioinformatics analysis with multi-agent systems. Sci Rep. 2025;15:39036. doi:10.1038/s41598-025-25919-z.

[9] Alam K, Roy B. From Prompt to Pipeline: large language models for scientific workflow development in bioinformatics. arXiv:2507.20122; 2025. https://arxiv.org/abs/2507.20122

[10] Cynthia ST, Roy B. Towards LLM-powered task-aware retrieval of scientific workflows for Galaxy. arXiv:2511.01757; 2025. https://arxiv.org/abs/2511.01757

[11] Meta. Llama 4: a new crop of flagship AI models. TechCrunch, April 5, 2025. https://techcrunch.com/2025/04/05/meta-releases-llama-4-a-new-crop-of-flagship-ai-models/

[12] Alibaba (Qwen team). Qwen3.6 family. GitHub. https://github.com/QwenLM/Qwen3.6

[13] DeepSeek-AI. DeepSeek-V4 preview release notes. DeepSeek API documentation, April 24, 2026. https://api-docs.deepseek.com/news/news260424

[14] Mistral AI. Introducing Mistral 3. Mistral AI news, December 2, 2025. https://mistral.ai/news/mistral-3

[15] Google. Introducing Gemma 4. Google blog, April 2, 2026. https://blog.google/innovation-and-ai/technology/developers-tools/gemma-4/

[16] IBM Research. Granite 4.1 AI foundation models. April 30, 2026. https://research.ibm.com/blog/granite-4-1-ai-foundation-models

[17] Allen Institute for AI (Ai2). OLMo 3. November 20, 2025. https://allenai.org/blog/olmo3

[18] Microsoft. Welcome to the new Phi-4 models — Phi-4-mini and Phi-4-multimodal. Microsoft TechCommunity, Educator Developer Blog. https://techcommunity.microsoft.com/blog/educatordeveloperblog/welcome-to-the-new-phi-4-models—microsoft-phi-4-mini–phi-4-multimodal/4386037

[19] NVIDIA. NVIDIA debuts Nemotron 3 family of open models. NVIDIA Newsroom. https://nvidianews.nvidia.com/news/nvidiadebuts-nemotron-3-family-of-open-models

[20] Cohere. Models — Command A, Aya. Cohere docs. https://docs.cohere.com/docs/models

[21] BestValueGPU. Consumer NVIDIA RTX GPU price history and specifications (RTX 4060 Ti, 4090, 5060 Ti, 5070, 5090). https://bestvaluegpu.com/

[22] Apple. Mac Studio configurations and pricing. https://www.apple.com/mac-studio/specs/

[23] Thunder Compute. NVIDIA RTX Pro 6000 Blackwell pricing analysis. https://www.thundercompute.com/blog/nvidia-rtx-pro-6000-pricing

[24] Thunder Compute. AMD Instinct MI300X pricing. https://www.thundercompute.com/blog/amd-mi300x-pricing

[25] Jarvis Labs. NVIDIA H100 80 GB pricing guide. https://jarvislabs.ai/blog/h100-price

[26] Northflank. NVIDIA B200 cost analysis and cloud rental rates. https://northflank.com/blog/how-much-does-an-nvidia-b200-gpu-cost

[27] Nekrutenko A. Datasets for Galaxy Collection Operations Tutorial. Zenodo dataset, 2021. doi:10.5281/zenodo.5119008. https://zenodo.org/records/5119008

[28] Rebolledo-Jaramillo B, Su MS, Stoler N, McElhoe JA, Dickins B, Blankenberg D, Korneliussen TS, Chiaramonte F, Nielsen R, Holland MM, Paul IM, Nekrutenko A, Makova KD. Maternal age effect and severe germ-line bottleneck in the inheritance of human mitochondrial DNA. Proc Natl Acad Sci U S A. 2014;111(43):15474–15479. https://pubmed.ncbi.nlm.nih.gov/25313049/

[29] Nekrutenko A. iwc-workflows/haploid-variant-calling-wgs-pe (v0.1). Zenodo, March 24, 2025. doi:10.5281/zenodo.15078463. https://zenodo.org/records/15078463

[30] Maier W, Bray S, van den Beek M, Bouvier D, Coraor N, Miladi M, Singh B, De Argila JR, Baker D, Roach N, Gladman S, Coppens F, Martin DP, Lonie A, Grüning B, Kosakovsky Pond SL, Nekrutenko A. Ready-to-use public infrastructure for global SARS-CoV-2 monitoring. Nat Biotechnol. 2021;39(10):1178–1179. https://pubmed.ncbi.nlm.nih.gov/34588690/

[31] Mei H, Arbeithuber B, Cremona MA, DeGiorgio M, Nekrutenko A. A high-resolution view of adaptive event dynamics in a plasmid. Genome Biol Evol. 2019;11(10):3022–3034. https://pubmed.ncbi.nlm.nih.gov/31539047/

[32] Galaxy Project Intergalactic Utilities Commission (IUC). tools-iuc: community-curated Galaxy tool wrappers, commit 39e7456. GitHub. https://github.com/galaxyproject/tools-iuc

